# Study replication: Shape discrimination in a conditioning procedure on the jumping spider *Phidippus regius*

**DOI:** 10.1101/2023.04.25.538063

**Authors:** Eleonora Mannino, Lucia Regolin, Enzo Moretto, Massimo De Agrò

**Affiliations:** Department of General Psychology, University of Padua, Italy; Esapolis’ Living Insects Museum of the Padua Province, Padua, Italy; Butterfly Arc Ltd., Padua, Italy; Biorobotics Department, Sant’Anna School of Advanced Studies, Pisa, Italy

**Keywords:** Amodal completion, Gestalt, Vision, Learning, Invertebrates

## Abstract

Jumping spiders possess a unique visual system, split into 8 different eyes and divided into two fully independent visual pathways. This peculiar organization begs the question of how visual information is processed, and whether the classically recognized gestalt rules of perception hold true. In a previous experiment, we tested the ability of jumping spiders to associate a geometrical shape with a reward (sucrose solution), and then to generalize the learned association to a partially occluded version of it. The occluded shape was presented together with a broken version of the same shape. The former should be perceived as a whole shape only in the case the animals, like humans, are able of amodally complete an object partly hidden by an occluder; otherwise, the two shapes would be perceived as identical. There, the spiders learned the association but failed to generalize. Here, we present a replication of the experiment, with an increased number of subjects, a DeepLabCut-based scoring procedure, and an improved statistical analysis. The results of the experiment follow closely the direction of the effects observed in the previous work but fail to raise to significance. We discuss the importance of study replication, and we especially highlight the use of automated scoring procedures to maximize objectivity in behavioural studies.

## Introduction

The ability to recognize and categorize individual objects is critical for survival for many animals. Although this is true for all the senses, vision has been the primary focus of such processes, and how it allows many creatures to pursue all major life activities including feeding, predator avoidance, social interactions, and sexual behavior (Lazareva et al., 2012). The visual scene is however constituted by a collection of seemingly unconnected elements and stimuli, that need to be organized by the brain into meaningful units which are then recognized as objects. Gestalt psychology describes a set of rules according to which our minds organize and interpret visual data (Vezzani et al., 2012; Wertheimer, 1923), under the assumption that the whole is greater than its parts. In other words, we do not simply focus on every separate component of the visual scene; instead, we aggregate multiple elements according to some similarities and perceive their global configuration. Such configuration is perceptually different from the array of elements composing it, in spite of the fact that the local information is identical in both cases.

An immediate application of such a system is apparent when considering the commonality of partially occluded figures. In real-world, objects present in the environment can often overlap: the ones closer to the observer can partially cover smaller or larger portions of the ones located further away. Nevertheless, we remain capable of recognizing the single fragments as a single object, as the visible parts are sufficient for completing the contour. Amodal completion is the ability of perceiving completed objects even in cases of partial occlusion (Kanizsa, 1979; Michotte, 1963), integrating fragmentary visual information, such as object shape, size, position, and numerosity from spatially segregated edges, and thus allowing to recognize an object despite the absence of any view of its covered parts. Gestalt psychology has described a set of perceptual rules, that define which fragments get collected amodally into a single objects, and which are considered separate (Vezzani et al., 2012). This mechanism has been widely studied in many species, including baboons (Deruelle et al., 2000), capuchin monkeys (Fujita & Giersch, 2005), chimpanzees (O’Connell & Dunbar, 2005; Sato et al., 1997), young chicks (Regolin & Vallortigara, 1995), pigeons (Nagasaka & Wasserman, 2008), mice (Kanizsa et al., 1993), fish (Sovrano et al., 2022) and overall seemingly ubiquitous across all vertebrates taxa. This is probably due to the overarching need of for animals to recognize meaningful objects with from partial incomplete information, this would be essential for visually guided foraging or for identification of conspecifics and predators. However, even if this aggregation approach to object recognition seems to be quite successful, using the global configuration of the stimuli is not the only solution. Some animals do not seem to need the global information (Sato et al., 1997; van Hateren et al., 1990), in favour of the analysis of specific local features (Kanizsa et al., 1993; Sovrano & Bisazza, 2008), like peculiar colours or marking that can trigger recognition without the need for information on the global configuration.

Even though the organization of their perceptual system is profoundly different from ours, the study of vision has also widely extended to invertebrates. For example, the amodal completion mechanism has been observed to be present even in cuttlefishes (Lin & Chiao, 2017). Among arthropods, bees can for example use different characteristics to detect and recognize flowers, such as shape (Lehrer, 1999), color or luminance contrast (Chittka & Menzel, 1992; Ibarra et al., 2000), pattern orientation (van Hateren et al., 1990), and symmetry (Giurfa et al., 1996). All of these examples require the perception of the global configuration (i.e. the flower). On the other hand, social wasps can distinguish nest-mates based on subtle details such as facial markings (Sheehan & Tibbetts, 2011), highlighting the importance of local cues. Similarly, spiders can discriminate between types of prey using visual cues (Harland & Jackson, 2000), while mantids use a ‘perceptual envelope’ that include size, contrast with the background, and location in the visual field to classify a stimulus as potential prey (Prete, 2004), all without requiring the “full picture”.

Among arthropods, salticids display some of the richest visually guided behaviours (De Agrò, 2020; De Agrò et al., 2021; Herberstein, 2011; Jackson & Harland, 2009; Rößler et al., 2022; Shamble et al., 2017). Their highly developed visual system is composed of 4 specialized and anatomically distinct pairs of eyes. The forward-facing pair of ‘anterior-medial’ eyes is specialized in binocular and colour vision (Lim & Li, 2006; Nakamura & Yamashita, 2000; Zurek et al., 2015), possesses a high visual acuity and is characterized by a narrow (< 5°) field of view (Land, 1985; Land & Nilsson, 2012). The other three pairs of eyes are positioned around the cephalothorax, the ‘posterior-lateral’, ‘posterior-medial’, and ‘anterior-lateral’ pairs and grant the spider an almost 360° field of view (Land, 1985). These eyes are monochrome and have a lower visual acuity, but can detect motion (Land, 1971, 1972a, 1972b; Zurek et al., 2010; Zurek & Nelson, 2012) thanks to the wide visual field and fast response time. Given these characteristics, jumping spiders have quickly attracted the interest of scientist, wondering whether Gestalt principles still hold true in a visual system organized to separately detect different elements of the visual scene. For example, Dolev and Nelson (2014) found that the jumping spider *Evarcha culicivora*, specialized in hunting mosquitoes, can recognize its preferred prey based on simple “stick figures” of it. Crucially, the spiders seemed to use the relative orientation of the body elements (abdomen, legs) relatively to each other, suggesting the need of representing the whole-body to allow recognition. However, recognition remained even if the position of the elements was scrambled, suggesting a focus on the local characteristics. In another experiment, Rößler et. al (2022) presented *Salticus scenicus* with 3d models of bigger jumping spiders, which caused the subjects to run away after an initial freezing. When presented with an equally sized 3d blob, *Salticus scenicus* had no reaction. Curiously, if the spiders were presented with a 3d blob with spider eyes pasted on it, the subjects had a mixed reaction, with some of them freezing and escaping, while others did not respond. This suggest that while the local feature of the eyes is relevant, this is not always sufficient, requiring the whole figure to trigger recognition. As it stands, the validity of gestalt principles in jumping spiders remains unclear.

In a previous study (De Agrò et al., 2017), we trained the jumping spider *Phidippus regius* to discriminate among two geometrical shapes, a circle and a cross. Afterwards, we asked the animals to choose between two versions of the rewarded shape: one partially occluded, and one cut in sections. If jumping spiders were able to complete amodally the missing parts, we should have observed a generalization of the learned association towards the occluded version. Conditioning was effective, as spiders were able to learn the discrimination task. However, they did not generalize the association to the illusory stimulus, suggesting that spiders’ visual system may not distinguish among the two configurations present at test. It may well be that jumping spiders do not require amodal completion. Nevertheless, before expanding on this hypothesis, some considerations need to be made on the shortcomings of our previous study. First, the experiment included only 18 subjects, fact that coupled with the low-response rate of the experiment significantly lowered the statistical power of the analysis. Even more crucially, spiders’ choices were scored manually by the experimenters, that had to judge from the videos whether the spiders consumed the presented reward. This left the entirety of decision making to the experimenters themselves concerning which trial had to be included in the analysis. This can introduce important subconscious biases in the scoring: even though a double-blind procedure was implemented, an experienced scorer could pick up subtle elements of the spiders’ behaviours suggesting the experimental condition, as well as whether the food contacted by the spider was a reward or not. Especially given the low number of subjects, the experimenter’s decisions of counting or not a behaviour as choice in a handful of trials had a strong weight on the final results (De Agrò, 2019).

In the current study, we replicated fully the experiment presented in De Agrò et al. (2017). However, we increased the number of subjects, as well as implementing an automated scoring procedure, based on the pose-estimation software DeepLabCut (Mathis et al., 2018). This was done to thoroughly remove the experimenters from the decision making process of data inclusion and video scoring, so as to guarantee maximum objectivity.

## Materials and Methods

A new group of jumping spiders underwent exactly the same experimental procedure described in De Agrò et al. (2017). However, the scoring and analysis procedure are completely new.

### Subjects

Thirty between juveniles and adults *Phidippus regius* were born in captivity and were kept individually in transparent plastic boxes (7×16×6 cm); rearing boxes were enriched with a portion of a cardboard egg container to build nests and hide, and a water dispenser that was topped-up as needed. During their life, spiders were fed ad libitum with *Tenebrio molitor* larvae. This feeding routine was interrupted 3 days before the start of the testing period to ensure motivation. Furthermore, they were received one *Tenebrio molitor* larvae every week during the breaks of the tasks. Lighting in the laboratory consisted of neon lamps with natural light (5000 kelvin colour temperature, 36 watts, 3350 lumens) on a 12h light/dark cycle with lights on at 08:00h. Subjects were well laboratory acclimated before testing and they had no previous experience with the types of stimuli presented. Tests were carried out between 09:00 and 18:00h.

### Apparatus

Trials were performed in plastic boxes of the same size as the housing ones. The walls of the boxes were lined from the outside with white paper to preventing subject’ viewing other spiders and the lab during tests, while the lid remained transparent to allow the light to pass through and to let the camera to record (Figure 1A). Two L-shaped white plastic blocks (Figure 1B) were set in each box and two red-colored drops (≈40 μl) were placed behind them so that spiders could not see them from their starting visual perspective. As in the previous study, the vertical wall of the white blocks showed either a ‘O’ or an ‘X’, both colored in black and matched for total area (7 mm^2^). Each drop contained sugar (20%) (as positive reward) or citric acid (25%) (as punishment) and they were visually identical; however, since sugar absorbs ultra-violet light (De Voe, 1975), spiders could have perceived the color of the drops differently, creating the need for the vertical plastic screen hiding the drop from sight from the perspective of the starting position.

**Figure 1.**
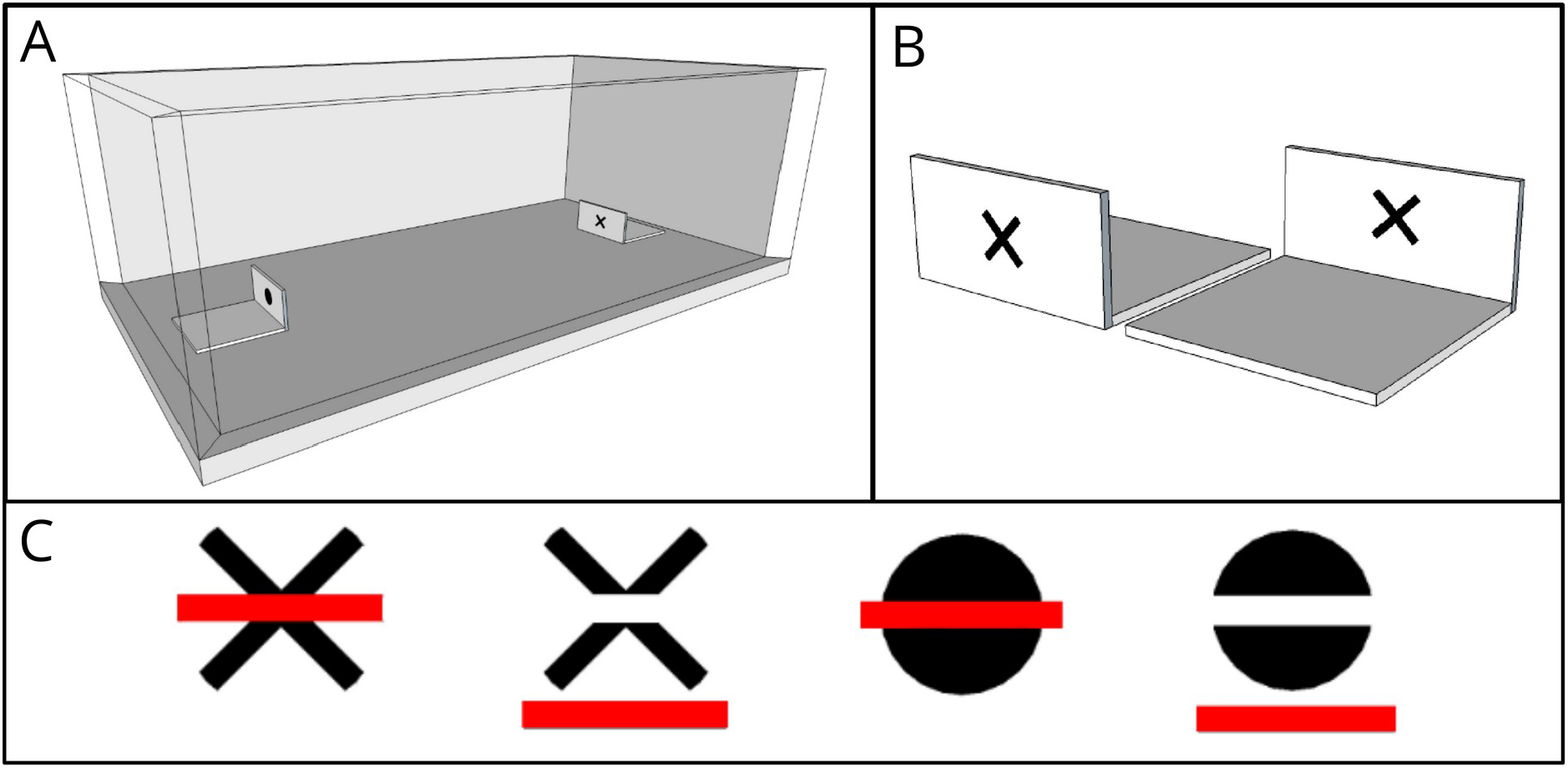
A: Experimental apparatus (7×16×6 cm) with the two platforms, X and O, located at both the short side ends of the arena. Each spider was placed in the centre of the arena and had 45 minutes to freely explore and choose to approach either figure. B: Example of the “X” platform where the drop (40 μl) was placed; front and back views. C: Stimuli employed in the amodal completion test. From the left, spiders trained on the “X” shape were exposed to the first vs second stimuli, while spiders trained on the “O” shape were exposed to the third vs fourth stimuli. Each pair of stimuli are identical as for the local features of the black area, but differ in their global configuration (to the human eye) (figures from De Agrò et al., 2017).

### Design and Procedure

Each individual was subjected to a training phase and a testing phase. To ensure a high level of motivation and maximize responsiveness, spiders were starved for 7 days prior to testing. The day before testing, spiders were placed in new empty boxes that were identical as the housing ones, where they were kept for the entire experiment duration.

For each of 6 days, every subject performed 3 trials, with a 1 hour long interval among trials to avoid excessive stress. At the start of each trial, spiders were carefully placed in the center of the box, while the L-shaped blocks with the red-colored drops were positioned at the short sides of the box; this way, they could easily see the figures from within their visual perspective form the starting position, but not the drops. Once in place, the spider was given 45 min to freely move around the box and it had free access to the water drops. The shapes were randomly assigned, and we kept the combination of figure and sugar or citric acid constant for each subject. To ensure that spiders learned to associate the figure with the drop flavour, the location was changed trial by trial.

The testing procedure lasted for 2 weeks, and it took place 2 days after the training phase. Each subject accomplished 18 trials, divided between 12 ‘rewarded trials’ (RT), 3 ‘unrewarded shape’ trials (US) and 3 ‘ unrewarded illusion’ trials (UI). The RT were 2/3 of these 18 trials, and the same stimuli of the training phase were used. These were included to maintain the supposed learned association. The remaining amount of trial were split between US, where the stimuli were presented without the reward/punishment, and UI, where the spiders were tested with a completely new stimulus still without the presence of reward. In this last case, the stimuli presented were a cut version and an occluded version of the same shape the spider was trained on. We used red bar with a different luminance so that to be able to distinguish the stimuli, spiders were expected to perceive the whole shape behind the stripe, while considering novel the cut version (Figure 1C). This way, we tested whether the spiders relied on amodal completion or not.

### Scoring and data analysis

To record the experiment, we placed 4 boxes near each other, 60cm below a raspberry pi equipped with raspberry pi digital cameras (V1.3). Each camera recorded at 5fps and 1080p resolution. Using a total of 4 identical setup, we were able to record 16 spiders at once.

Afterwards, we used the python software DeepLabCut (Mathis et al., 2018) to train a neural network capable of recognizing 4 points on the spiders’ bodies (position of the two ALEs, pedicel and spinneret), as well as the position of the two drops. We then fed all the recorded videos to the trained network, that outputted the frame-by-frame coordinates of the trained points. Subsequently, using a custom script written in python (Van Rossum & Drake, 2009), using the packages NumPy (Oliphant, 2006; van der Walt et al., 2011), SciPy (Virtanen et al., 2020), and pandas (Reback et al., 2020), we recorded all the events where the two ALEs points entered in contact with the drop (defined by the distance from the trained drop point). These were considered potential choice events. All these occurrences were then watched by an experimenter, that manually removed false positives. This was needed mostly because DeepLabCut was unable to discriminate spiders walking on the box lid and spiders walking on the ground. As such, it recorded as choice events many occurrences where the animals simply walked on the ceiling above the drops. In this phase, the experimenter removed also other false positives, like event where the animal didn’t cross the selection area, or just sprinted across. Crucially, in order to minimize bias, the videos were prompted to the experimenter without an indication of the video condition, subject or date. All the remaining events were kept for the analysis. For every spider, we recorded the number of visits to each drop per each trial. Many spiders only contacted one of the two drops for the whole trial duration: in those cases, we also recorded a binomial preference, with 1 being only contacts with the rewarded shape, and 0 being only contacts with the punished one.

All the analysis were performed in R4.1.2 (R Core Team, 2021), using the libraries glmmTMB (Brooks et al., 2017), car (Fox & Weisberg, 2019), DHARMa (Hartig, 2020), emmeans (Lenth, 2018), ggplot2 (Wickham, 2016), reticulate (Ushey et al., 2021). Plots were generated using python and the packages seaborn (Waskom et al., 2017) and matplotlib (Hunter, 2007). We modelled the binomial choice using generalized mixed effect models with a binomial error structure. We included in the model both the trial type and trial number and included the trial type nested in subject identity as a random intercept. After evaluating the goodness of fit for the model, we observed the effect of the predictors using an analysis of deviance, and subsequently performed Bonferroni-corrected post-hoc analysis on factors that had a significant effect. We performed a second analysis on the number of visits to each option in each trial, using a generalized mixed effect model with a Poisson error structure. We included in the model the visit value (rewarded or unrewarded shape), the trial number and the trial type. We again included the trial type nested in subject identity as a random intercept. After again evaluating the goodness of fit and observing the effect with the analysis of deviance, we performed Bonferroni-corrected post-hoc analysis on relevant factors.

## Results

Raw data for the experiment is available in supplement 1. For the full analysis and script, see supplement 2. Only the most relevant results are reported below. All the tests were performed on week 2 and 3 of the experiment, as week 1 only included training trials. We observed a generally low response rate: spiders contacted any one of the two drops in only 7.4% of the RT trials, 9.1% of the US trials and 17.8% of the UI ones. The latter appeared to be a significantly higher proportion in respect to RT (post-hoc, Bonferroni corrected, odds. Ratio=2.727, SE=0.725, DF=1413, t=3.771, p=0.0005).

Considering the binomial choice, we observed a significant effect of trial number (GLMM Analysis of Deviance, χ2=6.0311, DF=1, p=0.0140), but no effect of trial type (χ2=3.958, DF=2, p=0.18307), nor of the interaction between the two (χ2=0.9012, DF=2, p=0.63725). The post-hoc revealed that spiders chose significantly more often the correct shape in the Rewarded trials (post-hoc Bonferroni corrected, prob.=0.786, SE=0.058, DF=106, t=3.771, p=0.0008), but performed at chance level in the Unrewarded-shape trials (prob.=0.664, SE=0.113, DF=106, t=1.350, p=0.5394) and the Unrewarded-illusion trials (prob.=0.578, SE=0.095, DF=106, t=0.806, p=1) (Figure 2A). We also observed a non-significant increase in correct choices across subsequent trials, independently of trial type (trend=0.0542, SE=0.0277, DF=106, t=1.957, p=0.0530).

**Figure 2.**
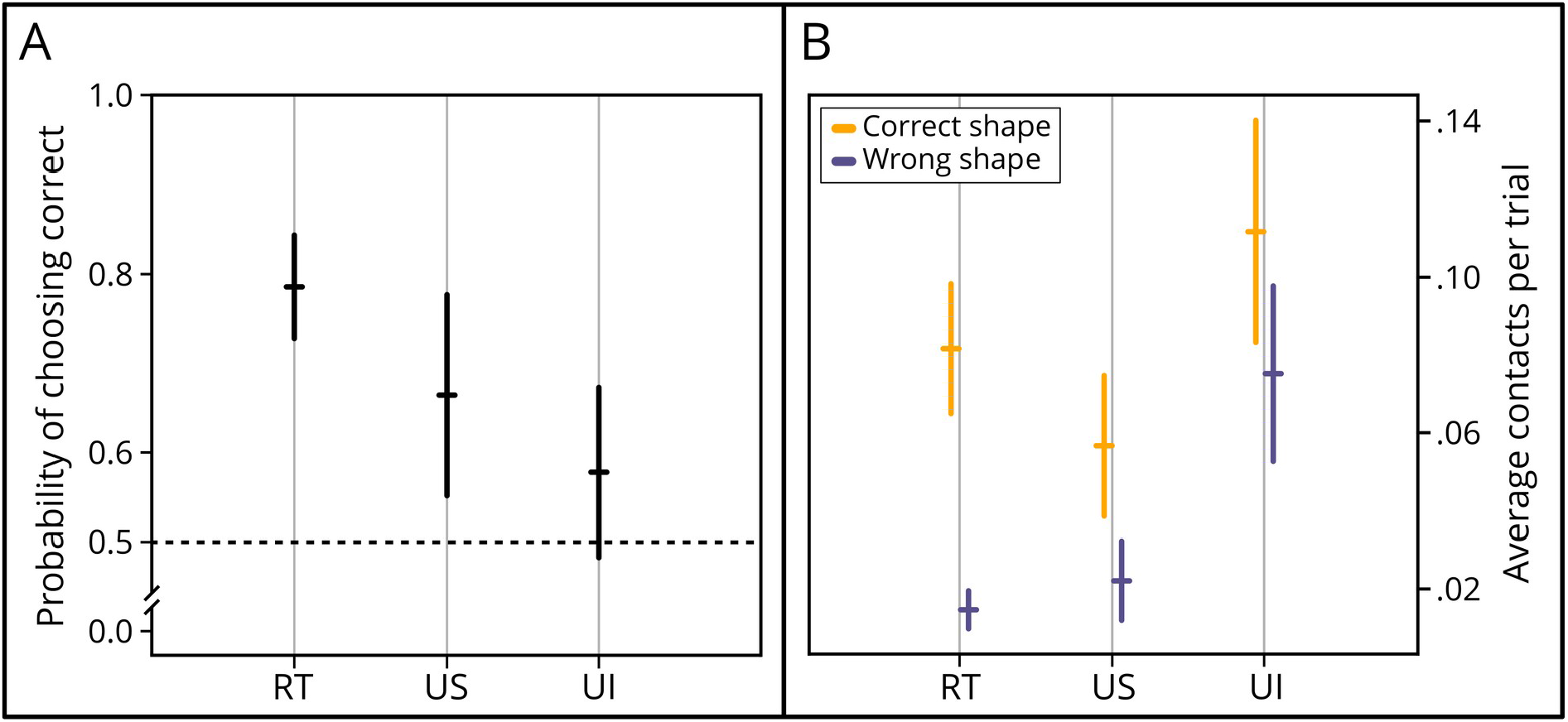
Results. A: Binomial choice. On the X axis is reported the trial type. On the Y axis, the probability of choosing the shape associated with sugar water. Spiders significantly preferred the rewarded shape in Rewarded Trials (RT, prob.=0.786, p=0.0008) but performed at chance level in the Unrewarded-shape trials (US, prob.=0.664, p=0.5394) and Unrewarded-illusion trials (UI, prob.=0.578, p=1). B: Contacts frequency to each drop. On the X axis is reported the trial type. On the Y axis, the average contacts per trial. Spiders significantly visited more often the correct option over the incorrect one in Rewarded trials (RT, ratio=5.57, p<0.0001), but they did not in Unrewarded-shape trials (US, ratio=2.56, p=0.1693) nor in Unrewarded-illusion trials (UI, ratio=1.49, S, p=0.5938).

The model performed using the number of visits to each option gave very similar results. We observed a significant effect of the visit value (whether it was to the rewarded or to the punished option. GLMM Analysis of Deviance, χ 2 =21.73, DF=1, p=0.0001). We also observed an effect of trial type (χ 2 =9.6217, DF=2, p=0.008), of trial number (χ 2 =6.3727, DF=1, p=0.0116), of the interaction between visit value and trial type (χ 2 =10.459, DF=2, p=0.0054) and of the interaction between visit value and trial number (χ 2 =12.1545, DF=1, p=0.0005). The post-hoc revealed that spiders visited significantly more often the correct option over the incorrect one in Rewarded trials (Post-hoc, Bonferroni corrected; ratio=5.57, SE=1.68, DF=2271, t=5.696, p<0.0001), but did not so in Unrewarded-shape trials (ratio=2.56, SE=1.265, DF=2271, t=1.909, p=0.1693) nor in Unrewarded-illusion trials (ratio=1.49, SE=0.456, DF=2271, t=1.288, p=0.5938). Regarding the trend across trials, we observed that the spiders contacted the wrong option less across trials for the Rewarded trials (trend=-0.082, SE=0.0241, DF=2271, t=-3.405, p=0.004) and for the Unrewarded-illusion trials (trend=-0.06, SE=0.0224, DF=2271, t=-2.684, p=0.0439) but not for the Unrewarded-shape trials (trend=-0.02101, SE=0.0419, DF=2271, t=-0.501, p=1) (Figure 2B). No trend was appreciable for the number of contacts with the correct option (trends: RT=-0.00817, US=-0.0049, UI=-0.0045, all p values > 0.05).

## Discussion

In this study, we replicated the study by De Agro’ et al. (2017), using a novel sample of spiders, the new sample being larger than the original one, and implementing a fully automated scoring procedure.

Here, the spiders performed significantly above chance level in the rewarded-shape condition, considering both the percentage of correct trials (78.6% of choices for sucrose solution drop) and the raw number of contacts to either drop (correct drop touched 5.57 times more than the incorrect one). This is consistent with what previously observed (De Agrò et al., 2017). The percentage of correct choices is far lower than the one observed in our previous study (96.8%). This is probably caused by the automatic scoring procedure: DeepLabCut unequivocally introduces a certain level of noise that can bring to false positives. On the other hand, incorrect events that may have been discarded during manual scoring due to experimenter bias are correctly included here. Regardless of the difference, this result confirms that the training procedure is an effective Heading 1motivational tool for jumping spiders: all individuals consistently avoided the citric acid solution, while approaching and consuming the sucrose one. It is crucial to point out however, that the performance in the rewarded-shape trials does not equate to a successful association between the reward and the conditioned stimuli. Indeed, the spiders could rely on other cues to avoid the citric acid solution, including drops scent or subtle color differences. These, even if inappreciable to the human eye, could be evident from the spiders’ *sensorium* (De Voe, 1975; Kienitz et al., 2022; Lim & Li, 2006).

In the unrewarded-shape condition, the spiders preferred the correct shape 66.4% of the times, but this preference resulted non-significant. Similarly, when looking at the raw number of visits, the spiders contacted the drop by the trained shape 2.56 times more, but, again, this did not appear significantly different from chance. This is in contrast with what we previously observed (De Agrò et al., 2017), where spiders did prefer the trained shape significantly more often (70.9% of the times, p=0.048) than the other. The results are however, in our opinion, not particularly dissimilar. Firstly, the correct choice probabilities in the two studies are comparable: both are around 70%, and both are lower than the correct choice probability in the rewarded-shape trials (78.6% here, 96.8% in the previous experiment). This may suggest that the spiders learned the task, but the overall low response rate decreased the statistical power of the analysis. On the other hand, it is possible that we previously committed a type I error, declaring a significant preference when there was none. It is impossible with the available data to tell which of the two options is correct. We believe that the best approach is to remain conservative, accept the null hypothesis and declare our previous results a type I error. As such, the spiders seemed to have failed to associate the punishing and rewarding solutions with the shapes.

In the unrewarded-illusion condition, the spiders preferred the occluded shape 57.8% of the times, a non-significant percentage. Similarly, when looking at the raw number of visits, the spiders contacted the drop by the occluded version of the shape 1.49 times more than the broken one, again not significantly. This is in accordance with what we previously observed, as in De Agro’ et al. (2017) the spiders chose the occluded shape exactly 50% of the times. In the absence of a significant result in the unrewarded-shape condition, we cannot make any inferences about the amodal completion skills of jumping spiders. Indeed, if the spiders did not associate the shape with a reward, both options in the shape-illusion condition would have no value, even if correctly perceived.

We believe that the DeepLabCut-based, automatic scoring procedure that we implemented is an enormous improvement to the procedure. For certain, this allowed us to increase enormously scoring objectivity by minimizing the human involvement. It should be noted however that this is not fully removed from the procedure, and it could never be. A human must train the network, a human-specified criterion for events detection had to be employed, and a human experimenter had to confirm the events. As such, biases could not have been removed, only decreased. Excluding the objectivity benefits, we also observed a significant decrease in scoring time and a virtual zeroing of person-hours invested. In this experiment we collected 1425 videos for a total of 1068 hours and 45 minutes of footage, that we would have previously re-watched fully to detect events. This limits greatly the scalability of the experiment, as adding a single subject increases the footage to score by 40 hours. DeepLabCut instead requires only an initial involvement to train the neural network of around 5 hours, independently of the number of subjects. Then, scoring time becomes negligible, and crucially requires no involvement by a human. The procedure comes with its own shortcomings. Mainly, it introduces a variable amount of noise in the data, that will in turn require a higher sample to be filtered out. This, coupled with the low response of this behavioral procedure, makes it very difficult to observe any significant results. We are persuaded that the benefits offered by automation greatly outweighs the costs. However, today a lot of established and reliable procedures are available to study both learning (De Agrò, 2020; Jakob & Long, 2016; Liedtke & Schneider, 2014; Long et al., 2015) and visual abilities (De Agrò et al., 2021; Ferrante et al., 2023; Moore et al., 2014; Peckmezian & Taylor, 2015; Zurek et al., 2010) in jumping spiders, making the one described here often a sub-optimal solution.

## Conclusions

In this paper, we trained jumping spiders in a shape discrimination task. Then, we asked whether they were capable of generalizing the learned association to occluded versions of the trained shape, demonstrating the presence of amodal completion abilities. The experiment was designed as a replication of De Agro’ et al. (2017). The results of this replica seemed similar to the one observed in the original study, but ultimately failed to rise to significance, leaving an open question on whether amodal completion is present in these animals. We hope that future research will aim at closing this knowledge gap, as understanding whether Gestalt principles hold true in a modular visual system could bring surprising new ideas. Indeed, spiders do live in a world full of occluding elements, mimetic predators and preys, and varied backgrounds. As such the ability to perceive complete objects in this complicated environment is paramount to their survival as well. If truly jumping spiders do not rely on the mechanism of amodal completion, we will have to assume that they apply a yet unknown different solution to the problem. We believe that such a solution may be found in the modular nature of these animals’ visual system, where the mismatches and overlaps of information coming from different eyes may inform figure segmentation, without the need for additional perceptual processes.

## Supplementary Materials

S1: raw data. S2: full analysis script.

## Data Availability Statement

Raw data is included as a supplemental material

## Author Contributions

Conceptualization, MDA, LR and EMo; Methodology, MDA, LR and EMo; Software, MDA; Validation, EMa, MDA, LR and EMo; Formal Analysis, MDA; Investigation, EMa and MDA; Resources, Emo; Data Curation, MDA; Writing – Original Draft Preparation, EMa; Writing – Review & Editing, EMa, MDA, LR; Visualization, MDA; Supervision, MDA, LR, EMo; Project Administration, MDA.

