## Supplemental Material 02 for "Study replication: Shape discrimination in a conditioning procedure on the jumping spider *Phidippus regius*"

S2 Scripts - Analysis


### S2 Scripts - Analysis

Abstract

This supplement provides the entire R script and output of the
statistical analysis we performed and figures produced, in their
original form. It is presented in the spirit of open and transparent
science, but has not been carefully curated.

### Setup

#### Prepare R environment

```
library(glmmTMB) #for mixed models
library(car) #for anova on mixed models
library(DHARMa) #for goodness of fit of the model
library(emmeans) #for post hoc
library(ggplot2) #to plot
library(reticulate)
```

#### Prepare Python environment

```
import pandas as pd
import matplotlib.pyplot as plt
import numpy as np
```

#### Load data

```
all <- read.csv(paste0(path, 'S1_raw_data.csv'))
all$date <- as.factor(all$date)
all$subj <- as.factor(all$subj)
all$corrshape <- as.factor(all$corrshape)
all$week <- as.factor(all$week)
all$type <- as.factor(all$type)
all$dropvalue <- as.factor(all$dropvalue)
all$orientation <- as.factor(all$orientation)
all$corrposition <- as.factor(all$corrposition)
```

```
path = r.path

all = pd.read_csv(path+'S1_raw_data.csv')
```

### Analysis

#### Preliminary analysis

Before proceeding with the analysis, we can check some overall
observations about the experiment.

##### Response rate

Before looking at preference, we can observe in how many trials the
subjects responded. We can also observe if this changes across time and
condition.

```
pm0 <- glmmTMB(diddrink~type*trialn+(type|subj), data = all, family = binomial())

simres <- simulateResiduals(pm0)
plot(simres, factor=TRUE)
```

```
## Warning in plot.window(...): parametro grafico "factor" non valido
```

```
## Warning in plot.xy(xy, type, ...): parametro grafico "factor" non valido
```

```
## Warning in title(...): parametro grafico "factor" non valido
```

```
Anova(pm0)
```

```
## Analysis of Deviance Table (Type II Wald chisquare tests)
## 
## Response: diddrink
##               Chisq Df Pr(>Chisq)   
## type        13.2141  2   0.001351 **
## trialn       2.3132  1   0.128277   
## type:trialn  4.0153  2   0.134305   
## ---
## Signif. codes:  0 '***' 0.001 '**' 0.01 '*' 0.05 '.' 0.1 ' ' 1
```

```
e <- emmeans(pm0, ~type, type='response')
```

```
## NOTE: Results may be misleading due to involvement in interactions
```

```
e
```

```
##  type   prob     SE  df asymp.LCL asymp.UCL
##  Ext  0.0909 0.0263 Inf    0.0509    0.1572
##  Ill  0.1782 0.0337 Inf    0.1214    0.2539
##  Rew  0.0737 0.0111 Inf    0.0546    0.0987
## 
## Confidence level used: 0.95 
## Intervals are back-transformed from the logit scale
```

```
pairs(e, adjust='bonferroni')
```

```
##  contrast  odds.ratio    SE  df null z.ratio p.value
##  Ext / Ill      0.461 0.169 Inf    1  -2.106  0.1055
##  Ext / Rew      1.257 0.430 Inf    1   0.670  1.0000
##  Ill / Rew      2.727 0.725 Inf    1   3.771  0.0005
## 
## P value adjustment: bonferroni method for 3 tests 
## Tests are performed on the log odds ratio scale
```

the response rate for “illusion” trials is double the one of the
others two. This doesn’t rise to significance vs the “Extinction”
trials, but it does vs “Reward”

It’s hard to find an explanation for this. I assume the reason is
that different stimuli elicit a new arousal in the spiders that are more
prone to explore. It is also crucial to point out however that reward
trials are double the other two combined. This may have very well
brought to a skewed distribution of lack of response.

#### Main analysis

##### Binomial data

Now to the main analysis, namely the preference. Week 0 is gonna be
removed as it is only training, and would confound the analysis

```
test <- subset(all, all$week != 0)

m0 <- glmmTMB(drinkcorrectbin~type*trialn+(type|subj), data = test, family = binomial())
```

```
## Warning in fitTMB(TMBStruc): Model convergence problem; non-positive-definite
## Hessian matrix. See vignette('troubleshooting')
```

```
## Warning in fitTMB(TMBStruc): Model convergence problem; false convergence (8).
## See vignette('troubleshooting')
```

```
simres <- simulateResiduals(m0)
plot(simres, factor=TRUE)
```

```
## Warning in plot.window(...): parametro grafico "factor" non valido
```

```
## Warning in plot.xy(xy, type, ...): parametro grafico "factor" non valido
```

```
## Warning in title(...): parametro grafico "factor" non valido
```

```
Anova(m0)
```

```
## Analysis of Deviance Table (Type II Wald chisquare tests)
## 
## Response: drinkcorrectbin
##               Chisq Df Pr(>Chisq)    
## type        15923.6  2  < 2.2e-16 ***
## trialn       5171.4  1  < 2.2e-16 ***
## type:trialn 10141.2  2  < 2.2e-16 ***
## ---
## Signif. codes:  0 '***' 0.001 '**' 0.01 '*' 0.05 '.' 0.1 ' ' 1
```

```
e <- emmeans(m0, ~type, type='response')
```

```
## NOTE: Results may be misleading due to involvement in interactions
```

```
e
```

```
## Warning in .qf.non0(object@V, x): Negative variance estimate obtained!
```

```
##  type     prob       SE  df asymp.LCL asymp.UCL
##  Ext  6.80e-06 0.000104 Inf     0.000     1.000
##  Ill  5.97e-01 0.101688 Inf     0.393     0.772
##  Rew  8.40e-01      NaN Inf       NaN       NaN
## 
## Confidence level used: 0.95 
## Intervals are back-transformed from the logit scale
```

```
test(e, adjust='bonferroni')
```

```
## Warning in .qf.non0(object@V, x): Negative variance estimate obtained!
```

```
##  type     prob       SE  df null z.ratio p.value
##  Ext  6.80e-06 0.000104 Inf  0.5  -0.773  1.0000
##  Ill  5.97e-01 0.101688 Inf  0.5   0.929  1.0000
##  Rew  8.40e-01      NaN Inf  0.5     NaN     NaN
## 
## P value adjustment: bonferroni method for 3 tests 
## Tests are performed on the logit scale
```

```
toplot <- as.data.frame(e)
```

```
## Warning in .qf.non0(object@V, x): Negative variance estimate obtained!
```

```
ggplot(toplot, aes(x=type, y=prob))+
        geom_point()+
        geom_errorbar(aes(ymin=prob-SE, ymax=prob+SE))+
        ylim(0,1)+
          geom_hline(yintercept = 0.5)
```

Something very strange is happening here. The model declares that
extinction trials have a 0% probabilty of correct, but this is not
significantly different from 50%? it is impossible. Something is
happening in the model. Will redo with type as random intercept rather
than slope.

```
m0 <- glmmTMB(drinkcorrectbin~type*trialn+(1|subj:type), data = test, family = binomial())

simres <- simulateResiduals(m0)
plot(simres, factor=TRUE)
```

```
## Warning in plot.window(...): parametro grafico "factor" non valido
```

```
## Warning in plot.xy(xy, type, ...): parametro grafico "factor" non valido
```

```
## Warning in title(...): parametro grafico "factor" non valido
```

```
Anova(m0)
```

```
## Analysis of Deviance Table (Type II Wald chisquare tests)
## 
## Response: drinkcorrectbin
##              Chisq Df Pr(>Chisq)  
## type        3.3958  2    0.18307  
## trialn      6.0311  1    0.01406 *
## type:trialn 0.9012  2    0.63725  
## ---
## Signif. codes:  0 '***' 0.001 '**' 0.01 '*' 0.05 '.' 0.1 ' ' 1
```

```
e <- emmeans(m0, ~type, type='response')
```

```
## NOTE: Results may be misleading due to involvement in interactions
```

```
e
```

```
##  type  prob    SE  df asymp.LCL asymp.UCL
##  Ext  0.664 0.113 Inf     0.424     0.842
##  Ill  0.578 0.095 Inf     0.390     0.746
##  Rew  0.786 0.058 Inf     0.651     0.878
## 
## Confidence level used: 0.95 
## Intervals are back-transformed from the logit scale
```

```
test(e, adjust='bonferroni')
```

```
##  type  prob    SE  df null z.ratio p.value
##  Ext  0.664 0.113 Inf  0.5   1.350  0.5307
##  Ill  0.578 0.095 Inf  0.5   0.806  1.0000
##  Rew  0.786 0.058 Inf  0.5   3.771  0.0005
## 
## P value adjustment: bonferroni method for 3 tests 
## Tests are performed on the logit scale
```

```
toplot <- as.data.frame(e)

et <- emtrends(m0, ~1, var = 'trialn')
et
```

```
##  1       trialn.trend     SE  df asymp.LCL asymp.UCL
##  overall       0.0542 0.0277 Inf -8.38e-05     0.109
## 
## Results are averaged over the levels of: type 
## Confidence level used: 0.95
```

```
test(et, adjust='bonferroni')
```

```
##  1       trialn.trend     SE  df z.ratio p.value
##  overall       0.0542 0.0277 Inf   1.957  0.0504
## 
## Results are averaged over the levels of: type
```

These results appear much more reasonable than before. Only the
reward condition is significantly higher than chance. Extinction trials
are at 66%, but not enough to raise over significance, probably due to
the low n of responses. The illusion condition is much nearer to chance
level (58%). Regardless, we will treat both as not significant, as it
appropriate. Similar story can be said about the effect of time: the
anoda signals an effect, but this does not go below .05 in the post hoc.
Marginally significant, choices for the correct shape increase by 5%
every trial.

```
ggplot(test, aes(x=trialn, y=drinkcorrectbin, color=type))+
        geom_jitter(width = 0.25, height = 0.05, alpha=0.5)+
        geom_smooth(method='glm')+
  geom_hline(yintercept = 0.5)
```

```
## `geom_smooth()` using formula = 'y ~ x'
```

```
## Warning: Removed 2171 rows containing non-finite values (`stat_smooth()`).
```

```
## Warning: Removed 2171 rows containing missing values (`geom_point()`).
```

```
ggplot(toplot, aes(x=type, y=prob))+
        geom_point()+
        geom_errorbar(aes(ymin=prob-SE, ymax=prob+SE))+
        ylim(0,1)+
          geom_hline(yintercept = 0.5)
```

##### Count data

in some trials, spiders contacted more than 1 drop. in this case the
binomial choice has been considered as NA, as preference cannot be
assessed if they tried both. We will repeat the analysis using all the
contacts, and modelling the actual number of selections per trial using
a poisson distribution.

```
m1 <- glmmTMB(drinkfreq~dropvalue*type*trialn+(type|subj), data = test, family = poisson())
```

```
## Warning in fitTMB(TMBStruc): Model convergence problem; singular convergence
## (7). See vignette('troubleshooting')
```

```
# does not converge like this. I suspect that type as a slope does not work in this dataset
m1 <- glmmTMB(drinkfreq~dropvalue*type*trialn+(1|subj:type), data = test, family = poisson())

simres <- simulateResiduals(m1)
plot(simres, factor=TRUE)
```

```
## DHARMa:testOutliers with type = binomial may have inflated Type I error rates for integer-valued distributions. To get a more exact result, it is recommended to re-run testOutliers with type = 'bootstrap'. See ?testOutliers for details
```

```
## Warning in plot.window(...): parametro grafico "factor" non valido
```

```
## Warning in plot.xy(xy, type, ...): parametro grafico "factor" non valido
```

```
## Warning in title(...): parametro grafico "factor" non valido
```

```
Anova(m1)
```

```
## Analysis of Deviance Table (Type II Wald chisquare tests)
## 
## Response: drinkfreq
##                         Chisq Df Pr(>Chisq)    
## dropvalue             21.7300  1  3.139e-06 ***
## type                   9.6217  2  0.0081409 ** 
## trialn                 6.3727  1  0.0115888 *  
## dropvalue:type        10.4589  2  0.0053566 ** 
## dropvalue:trialn      12.1545  1  0.0004897 ***
## type:trialn            0.9386  2  0.6254464    
## dropvalue:type:trialn  1.0851  2  0.5812704    
## ---
## Signif. codes:  0 '***' 0.001 '**' 0.01 '*' 0.05 '.' 0.1 ' ' 1
```

```
e <- emmeans(m1, ~type*dropvalue, type='response')
```

```
## NOTE: Results may be misleading due to involvement in interactions
```

```
e
```

```
##  type dropvalue   rate      SE  df asymp.LCL asymp.UCL
##  Ext  correct   0.0566 0.01805 Inf   0.03033    0.1058
##  Ill  correct   0.1117 0.02845 Inf   0.06780    0.1840
##  Rew  correct   0.0816 0.01669 Inf   0.05467    0.1218
##  Ext  wrong     0.0221 0.01014 Inf   0.00898    0.0543
##  Ill  wrong     0.0752 0.02255 Inf   0.04177    0.1353
##  Rew  wrong     0.0146 0.00487 Inf   0.00763    0.0281
## 
## Confidence level used: 0.95 
## Intervals are back-transformed from the log scale
```

```
contrast(e, list('RewCorrVsWrong'= c(0,0,1,0,0,-1),
                 'ExtCorrVsWrong'= c(1,0,0,-1,0,0),
                 'IllCorrVsWrong'= c(0,1,0,0,-1,0)), adjust = 'bonferroni')
```

```
##  contrast       ratio    SE  df null z.ratio p.value
##  RewCorrVsWrong  5.57 1.680 Inf    1   5.696  <.0001
##  ExtCorrVsWrong  2.56 1.265 Inf    1   1.909  0.1689
##  IllCorrVsWrong  1.49 0.456 Inf    1   1.288  0.5934
## 
## P value adjustment: bonferroni method for 3 tests 
## Tests are performed on the log scale
```

```
toplot2 <- as.data.frame(e)


et <- emtrends(m1, ~type*dropvalue, var = 'trialn')
et
```

```
##  type dropvalue trialn.trend     SE  df asymp.LCL asymp.UCL
##  Ext  correct       -0.00488 0.0261 Inf   -0.0561    0.0463
##  Ill  correct        0.00450 0.0179 Inf   -0.0306    0.0396
##  Rew  correct       -0.00817 0.0101 Inf   -0.0280    0.0117
##  Ext  wrong         -0.02101 0.0419 Inf   -0.1032    0.0612
##  Ill  wrong         -0.06010 0.0224 Inf   -0.1040   -0.0162
##  Rew  wrong         -0.08213 0.0241 Inf   -0.1294   -0.0349
## 
## Confidence level used: 0.95
```

```
test(et, adjust='bonferroni')
```

```
##  type dropvalue trialn.trend     SE  df z.ratio p.value
##  Ext  correct       -0.00488 0.0261 Inf  -0.187  1.0000
##  Ill  correct        0.00450 0.0179 Inf   0.251  1.0000
##  Rew  correct       -0.00817 0.0101 Inf  -0.807  1.0000
##  Ext  wrong         -0.02101 0.0419 Inf  -0.501  1.0000
##  Ill  wrong         -0.06010 0.0224 Inf  -2.684  0.0436
##  Rew  wrong         -0.08213 0.0241 Inf  -3.405  0.0040
## 
## P value adjustment: bonferroni method for 6 tests
```

very similar story here. In reward trials spiders contact more
correct drops over wrong ones. In extinction trials the ratio is not
significant, but the correct contacts are more than double the wrong
ones. Illusion trials are again more similar to each other.

```
ggplot(test, aes(x=trialn, y=drinkfreq, color=dropvalue))+
  facet_wrap(~type)+
        geom_jitter(width = 0.25, height = 0.05, alpha=0.5)+
        geom_smooth(method='glm')
```

```
## `geom_smooth()` using formula = 'y ~ x'
```

```
ggplot(toplot2, aes(x=type, y=rate, color=dropvalue))+
        geom_point(position = position_dodge(0.8))+
        geom_errorbar(aes(ymin=rate-SE, ymax=rate+SE),position = position_dodge(0.8))
```

The over trial graph reveals that really these frequencies remain
very low, and lines very flat. not very useful to look at the trial by
trial effect.

### Plots for manuscript

I am gonna now prepare the plots that will go in the publication.

```
 #will make it fancy on inkscape. for now no colors

binomres = r.toplot
countres = r.toplot2

fig, ax = plt.subplots()
plt.scatter(binomres['type'].values, binomres['prob'].values)
plt.vlines(binomres['type'].values,
           ymin=np.subtract(binomres['prob'].values,binomres['SE'].values),
           ymax=np.add(binomres['prob'].values,binomres['SE'].values))
plt.axhline(0.5)
plt.ylim((0,1))
```

```
## (0.0, 1.0)
```

```
fig.savefig(path+'binresults.svg', format='svg', dpi=1200)

plt.show()
```

```
fig, ax = plt.subplots()
plt.scatter(countres['type'].values, countres['rate'].values)
plt.vlines(countres['type'].values,
           ymin=np.subtract(countres['rate'].values,countres['SE'].values),
           ymax=np.add(countres['rate'].values,countres['SE'].values))
fig.savefig(path+'cntresults.svg', format='svg', dpi=1200)
plt.show()
```
